# Black sea cucumber (*Holothuria atra* Jaeger, 1833) rescues *Pseudomonas aeruginosa*-infected *Caenorhabditis elegans via* reduction of pathogen virulence factors and enhancement of host immunity

**DOI:** 10.1101/515783

**Authors:** Wan-Ting Lee, Boon-Khai Tan, Su-Anne Eng, Gan Chee Yuen, Kit Lam Chan, Yee Kwang Sim, Shaida Fariza Sulaiman, Alexander Chong Shu-Chien

## Abstract

A strategy to circumvent the problem of multidrug resistant pathogen is consumption of functional food rich in anti-infectives targeting bacterial virulence or host immunity. The black sea cucumber (*Holothuria atra*) is a tropical marine sea cucumer species traditionally consumed as remedy for many ailments. There is a paucity of knowledge the anti-infectives capacity of *H. atra* and the underlying mechanisms involved. The objectives of this study were to utilize the *Caenorhabditis elegans*-*P. aeruginosa* infection model to assess the anti-infective properties of *H. atra*. We first showed the capacity of a *H. atra* extract and fraction in promoting survival of *C. elegans* during a customarily lethal *P. aeruginosa* infection. The same chemical entities also attenuate the production of several *P. aeruginosa* virulence factors and biofilm. Treatment of infected transgenic *lys-7*::GFP worms with *H. atra* fraction restored the repressed expression of *lys-7*, a defense enzyme, indicating improved host immunity. QTOF-LCMS analysis revealed the presence of aspidospermatidine, an indole alkaloid and inosine. Collectively, our finding shows that *H. atra* confers survival advantage in *C. elegans* against *P. aeruginosa* infection through inhibition of pathogen virulence and eventually, the restitution of host *lys-7* expression.

## 1. Introduction

Increasing number of antibiotic resistance *P. aeruginosa*, an opportunistic pathogen are being reported globally.^1^ Attempts to overcome bacteria resistance through modification of existing drugs or drug discovery are often ineffectual.^2^ Collectively, these issues underscore the need for alternative therapies with different modes of targeting mechanism involving host immune response or pathogen virulence. In comparison to direct killing of pathogens, these paths are projected to invoke milder evolutionary pressure responsible for resistance development. Since nutrient status is a crucial contributing factor to immune fitness, functional food with immunomodulator characteristic may play a pivotal role in alleviation of infection.^3^

From the estimated 1000 varieties of sea cucumbers worldwide, around 20 species are valued as functional food.^4^ Sea cucumbers are reported to be effective remedy for asthma, hypertension, joint pain, sprains intestinal and urinary problems.^5, 6^ Correspondingly, studies on sea cucumber metabolites have reported multiple activities including anticancer, anticoagulant, anti-hypertension, anti-inflammatory, antimicrobial, antioxidant, antithrombotic, antihyperglycemic, anti-ageing and wound healing.^4, 7, 8^ These bioactivities are linked to a wide range of metabolites, such as triterpene glycosides, peptides, collagen, mucopolysaccharides, carotenoids, polyunsaturated fatty acids and phospholipids.^8-10^ Therefore, sea cucumber is an opportune strategy to discover novel and potent anti-infective entities. *Holothuria atra* Jaeger 1833 (Holothuridae), colloquially known as black sea cucumber or lollyfish, is one of the most commonly found sea cucumber species in the Indo-Pacific region, dwelling in seagrass beds and rocky reefs.^11^ Among the many medicinal properties linked with sea cucumbers, sea cucumbers are also traditionally consumed for its immune boosting properties.^5^ In tandem, metabolites with *in vitro* or *in vivo* immunomodulatory activities have been isolated from several sea cucumber species, making this taxonomy group an opportune organism for discovery of novel and potent anti-infective entities.^12-15^

The nematode *Caenorhabditis elegans* is a proven model for deciphering host–pathogen interactions due to its readiness for infection by numerous human pathogens.^16^ As a model organism, it is also supported by an array of tools useful for deciphering immunity related pathways.^17^ The *C. elegans* host-pathogen interaction is a proven powerful platform for discovery of anti-infectives and is frequently utilized to screen for novel anti-infectives against a myriad of pathogens.^18, 19^ An advantage in having both host and pathogen in a screening assay is early detection of undesirable toxic properties in hit candidates.^20^ In relation to food, *C. elegans* has been used as platform for deciphering various beneficial properties of food including longevity, tolerance, lipid metabolism and stress resistance.^21-23^

To date, the only published report of an anti-infective from marine invertebrates using the *C. elegan*s platform is from a tropical mollusk.^24^ Using the *C. elegans* host-pathogen platform, we investigated the anti-infective properties of *H. atra*. Overall, we demonstrate the capacity of an extract from this sea cucumber in rescuing worms from a lethal *P. aeruginosa* infection. The capacity of *H. atra* metabolites in impeding the production of well-characterized *P. aeruginosa* virulence factors and boosting an immune-related gene were also studied.

## 2. Results

### 2.1 *H. atra* promotes survival of *P. aeruginosa* PA14-infected *C. elegans* without direct bactericidal effect

The slow-killing assay showed methanol extract of *H. atra* causing a significantly higher percentage of worm survival (60.21± 6.7 %) at 48 hours after treatment, which is comparable to value obtained with curcumin (60.68 ± 8.6%), a known anti-infective against PA14 infection (Fig. 1). A lower percentage of survival (56 ± 7.6%) was obtained with the acetone extract. The bactericidal effect of the methanol extract on *P. aeruginosa* PA14 was determined. Extract concentrations ranging from 25 µg/mL to 1000 µg/mL did not reduce *P. aeruginosa* PA14 growth, as compared to streptomycin at 100 µg/mL (Supplementary 1). This extract was also non-detrimental towards the growth kinetic of *P. aeruginosa* PA14, as no significant difference was observed between *H. atra* and DMSO exposed bacteria population (Fig. 2).

**Figure 1.**
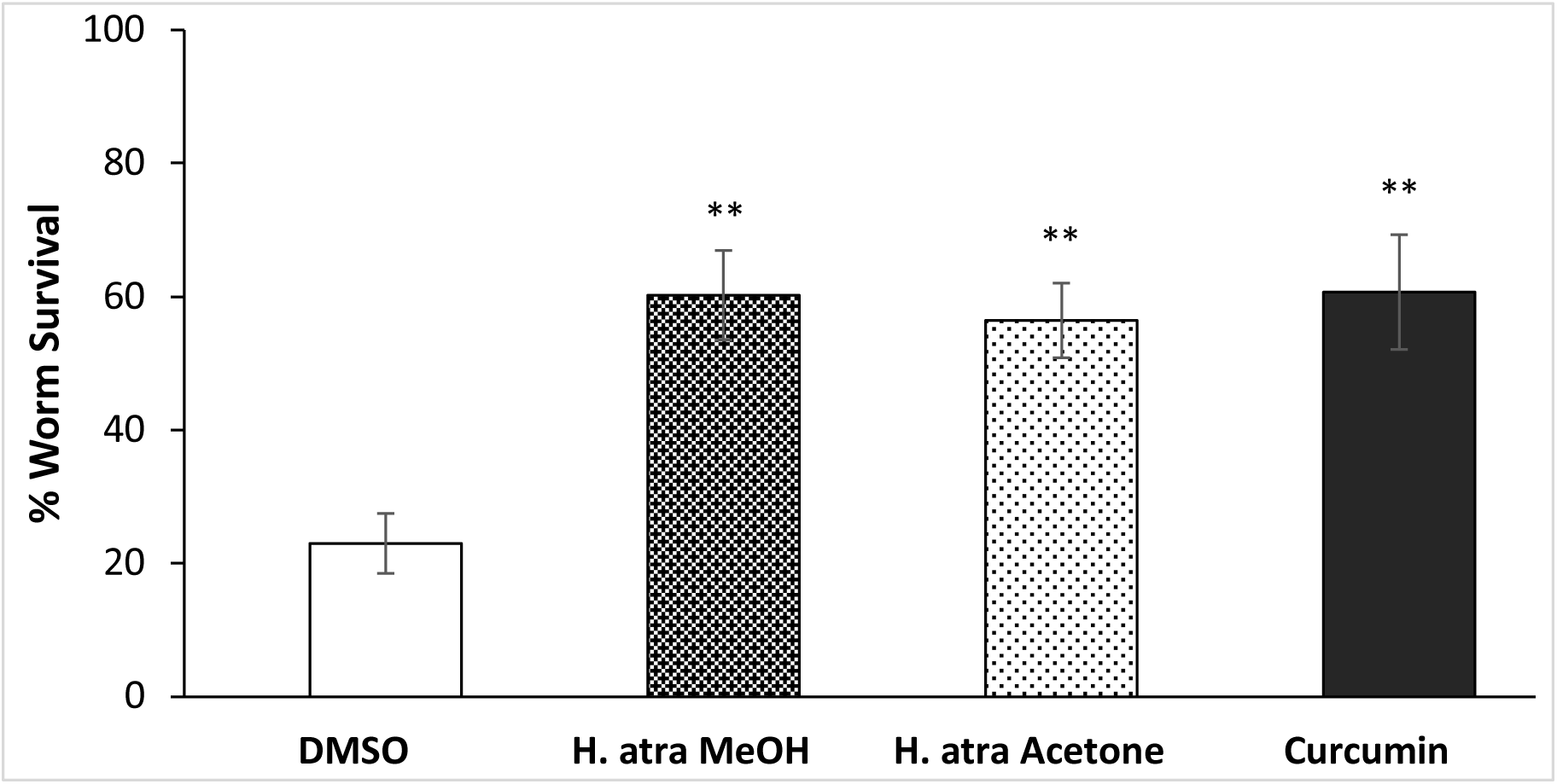
Improved survival of *P. aeruginosa* PA14-infected *C. elegans* at 48 hours after treatment with *Holothuria atra* extracts (200 µg/mL). The methanol extract showed the highest percentage of survival. Data were analyzed using Student’s *t*- test where ** denotes significance at *p*<0.01 when compared to 0.5% DMSO treatment. Curcumin was used as positive control.

**Figure 2.**
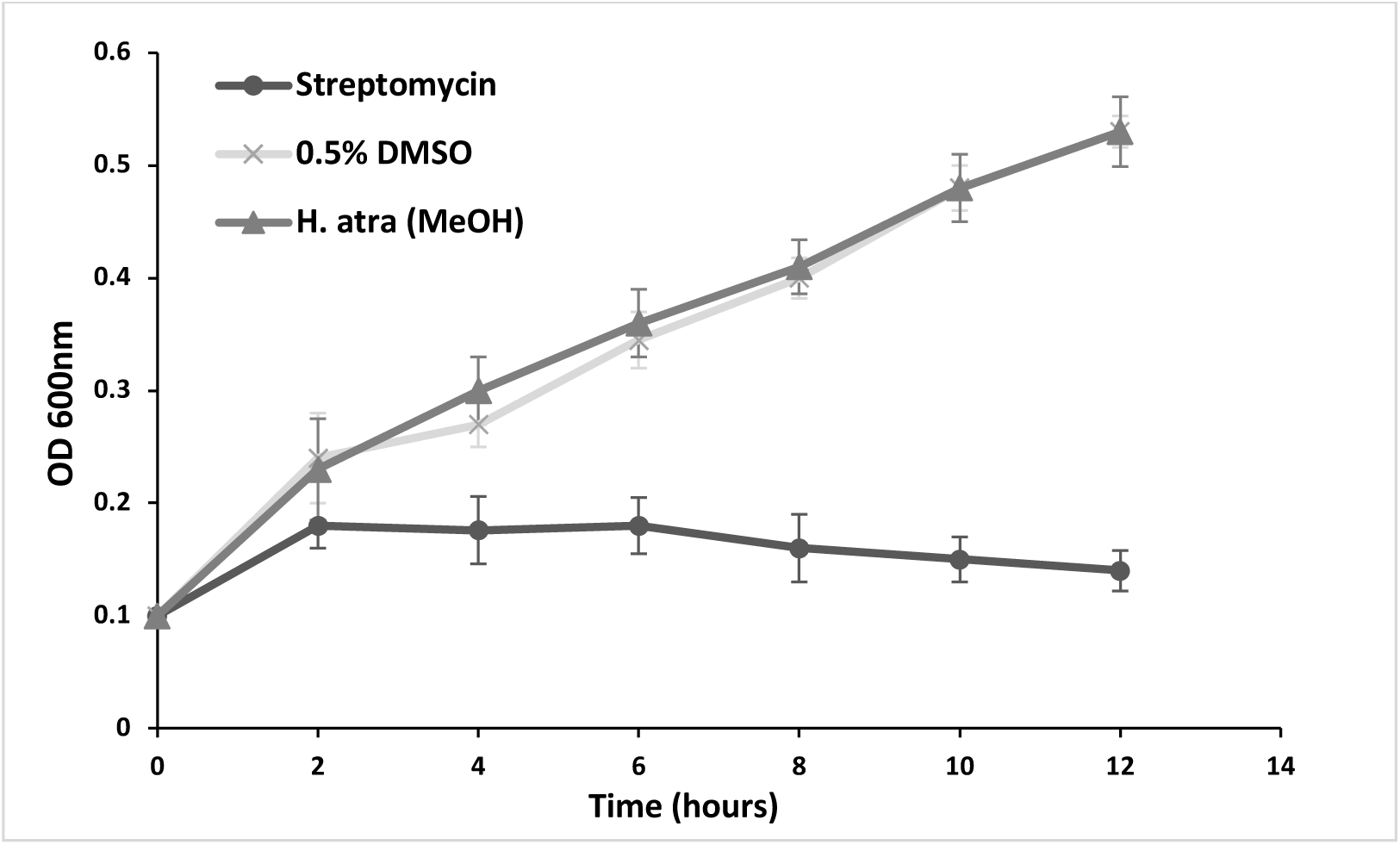
Hourly growth rate of *P. aeruginosa* PA14 in the presence of 200 µg/mL *Holothuria atra* methanolic extract. Growth was comparable to 0.5% DMSO treatment while streptomycin (100 µg/mL) reduced bacterial proliferation.

The lethality of *P. aeruginosa* PA14 infection in the slow killing assay involves the colonization of *C. elegans* intestinal tract.^25^ Therefore, we ascertain if the improved survival of worms treated with extract is an outcome of reduced worm feeding activities. We observed the worm pharyngeal pumping rate, a reliable indicator of bacterial feeding in this species.^24^ Results show no significant difference in feeding activities between untreated worms fed PA14 with those treated with either 0.5% DMSO or methanol *H. atra* extract, respectively (Fig. 3). Therefore, worms exposed to *H. atra* methanol extract still display normal feeding activities, which presumably leads to gut colonization by the ingested PA14 bacteria.

**Figure 3.**
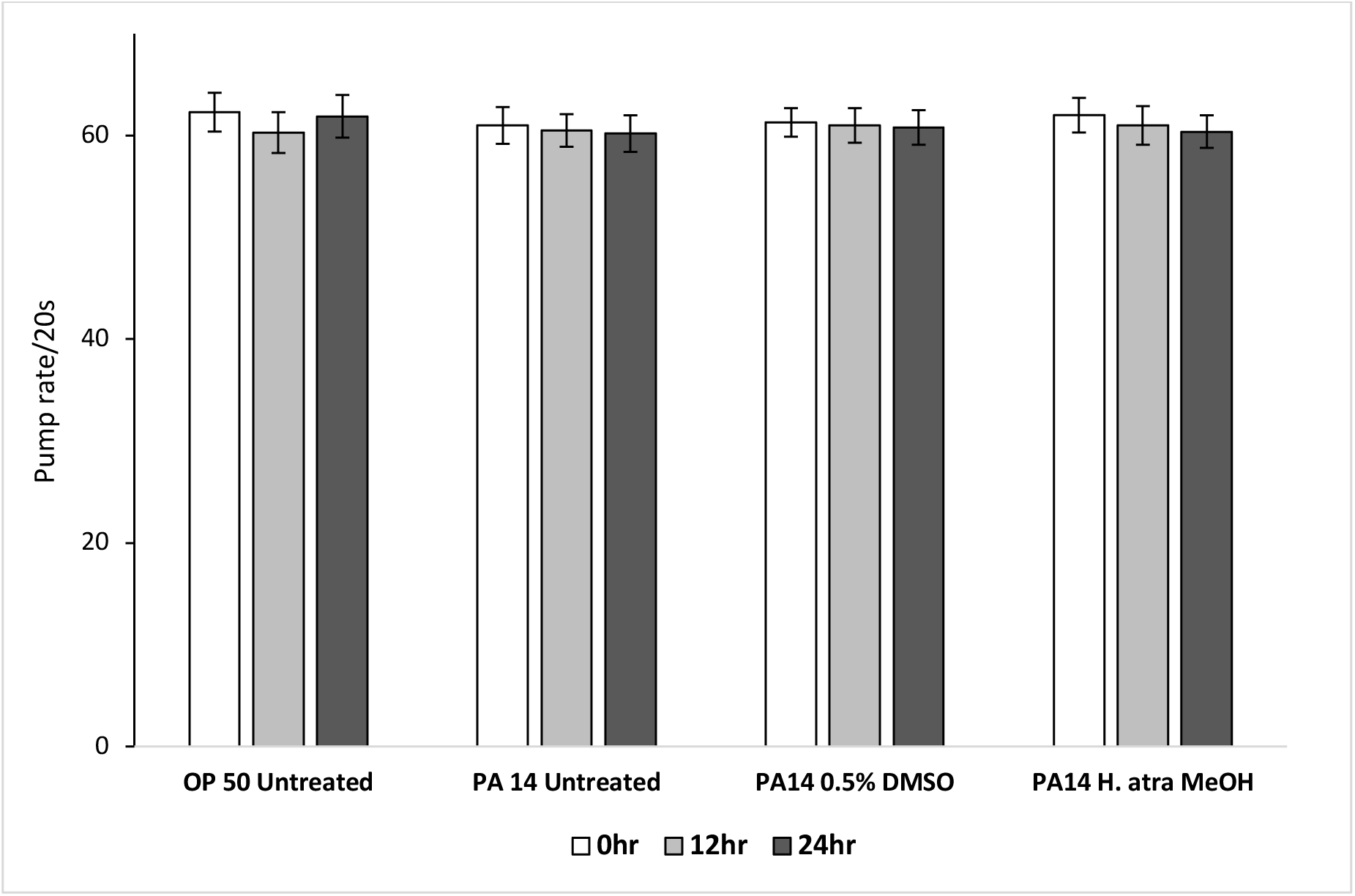
Pharyngeal pumping rate of *C. elegans* fed *E.coli* OP-50, *Pseudomonas aeruginosa* PA14 untreated, PA14 treated with 0.5% DMSO and PA14 treated with *Holuthuria atra* methanol extract at 200 µg/mL at 0, 12 or 24hr after treatment. Values were compared to 0.5% DMSO treatment for each respective hr after treatment using Student’s *t*-test at *p*<0.05 and *p*<0.01, respectively.

Column chromatography fractionation of the *H. atra* butanol partition resulted in 9 fractions (F1-F9). Based on the slow-killing assay, fractions F2 (55.37 ± 8.4%), F3 (64.05 ± 8.8%), F4 (52.75 ± 6.3%) and F9 (52.00 ± 8.0%) resulted in significantly higher worm survival as compared to 0.5% DMSO treatment (Fig. 4A). Dose dependent assay for fraction F3 showed that 200 µg/mL was the minimal concentration resulting in significantly highest percentage of worm survival, comparable to value obtained with curcumin (Fig. 4B). At higher concentrations, a decreasing trend of survival rates was observed. Fraction F3 also did not appear to perturb *P. aeruginosa* PA14 growth (Fig. 4C).

**Figure 4.**
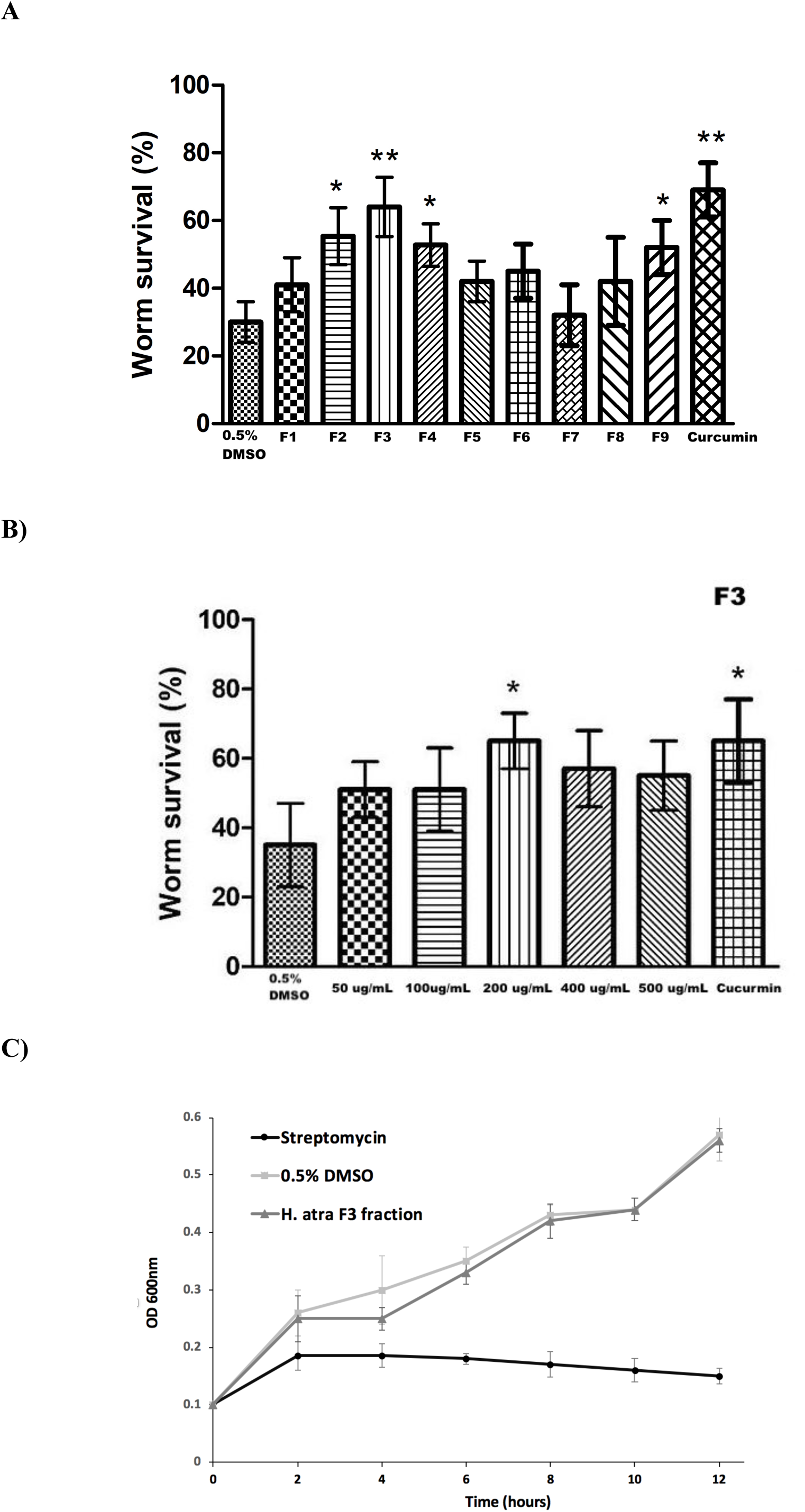
A) Survival of *P. aeruginosa* PA14-infected *C. elegans* at 48 hours after treatment with *Holothuria atra* fractions F1-F9. The F3 fraction showed the highest percentage of survival. Data were analyzed using Student’s *t*-test where * and ** denotes significance at *p*<0.05 and *p*<0.01 respectively when compared to 0.5% DMSO treatment. Curcumin, used as positive control, also prolonged worm survival significantly.B) Dose dependent response of *P. aeruginosa* PA14-infected *C. elegans* at 48 hours after treatment with *Holothuria atra* fraction F3. Data were analyzed using Student’s t-test where * at p<0.05 when compared to 0.5% DMSO treatment. Curcumin, as positive control, also prolonged worm survival significantly. C) Hourly growth rate of *P. aeruginosa* PA14 in the presence of 200 µg/mL *Holothuria atra* fraction F3. Growth was comparable to 0.5% DMSO treatment while streptomycin (100 µg/mL) reduced bacterial proliferation.

### 2.2 *H. atra* fraction inhibits the production of *P. aeruginosa* PA14 elastase, protease, pyocyanin and formation of biofilm

In comparison with 0.5% DMSO, treatment of *P. aeruginosa* PA14 with F3 significantly suppressed the production of elastase and protease at 24 hr after treatment (Fig. 5A). Formation of biofilm in microtiter plate was also attenuated with F3. As for pyocyanin, significant reduction in pyocyanin production was obtained at 6 and 12 hr after incubation, respectively, but not 24 hr. (Fig. 5B).

**Figure 5.**
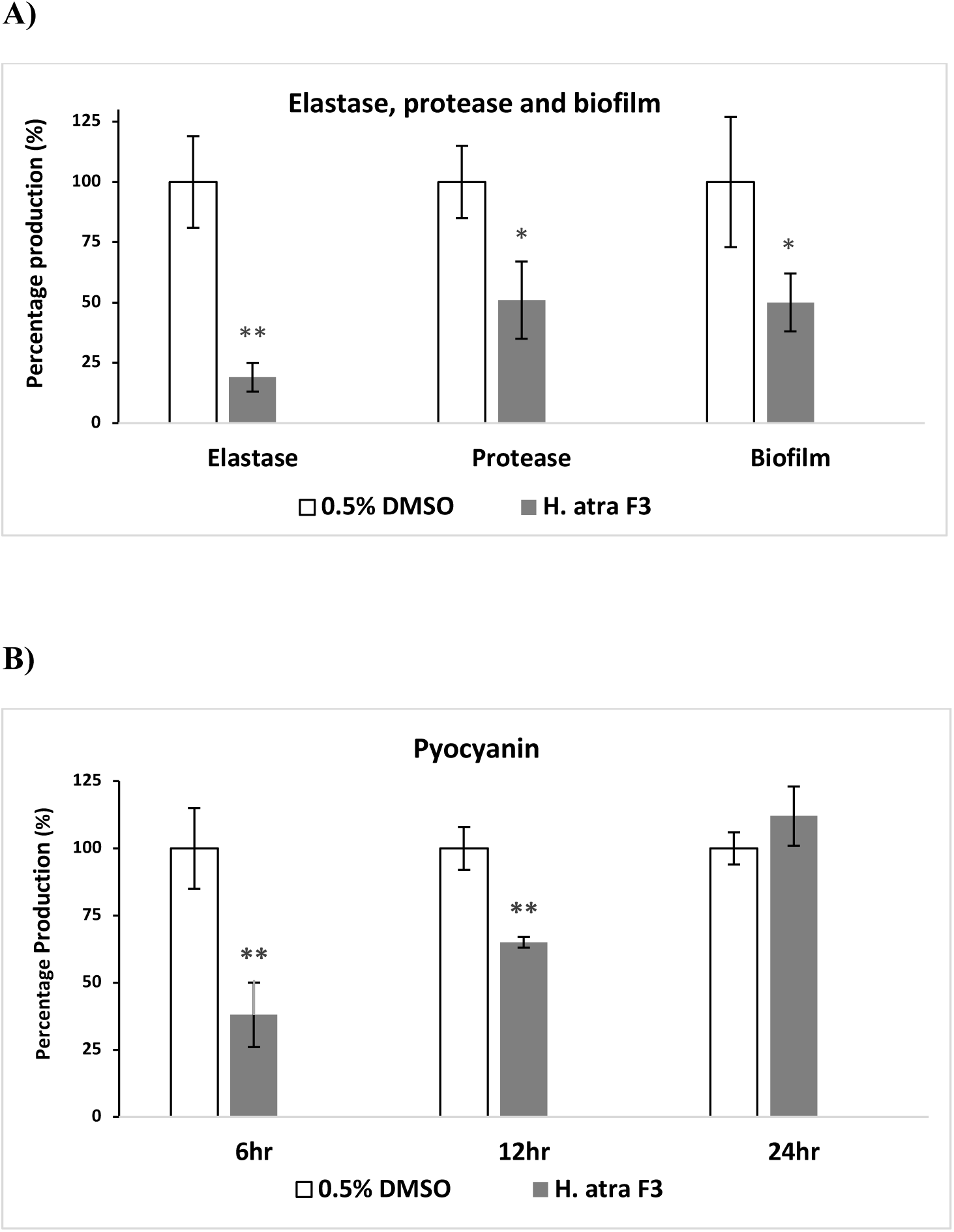
Effect of *Holothuria atra* F3 fraction on production of *P. aeruginosa* PA14 virulence factors (**A**) Reduced production elastase, protease and biofilm at 24 hr after exposure to F3 (**B**) Reduced pyocyanin production at 6 and 12 hr after exposure. Data were analyzed using Student’s *t*-test where * and ** denotes significance at *p*<0.05 and *p*<0.01 respectively when compared to 0.5% DMSO treatment.

### 2.3 *H. atra* induces the expression of *C. elegans* immune effector *lys-7* in PA14-infected worms

At 12 hr after infection, GFP intensity of *P. aeruginosa* PA14 infected worms was lower than uninfected worms, indicating diminished *lys-7* expression (Fig. 6A). Treatment with *H. atra* restored *lys-7* expression to a level comparable with uninfected worms. Correspondingly, qPCR analysis of *lys-7* in worms also showed higher expression levels in worms exposed to F3 at 12 or 24 hours after treatment (Fig. 6B). *H. atra* fraction *also* resulted in higher *lys-7* expression as compared to curcumin.

**Figure 6.**
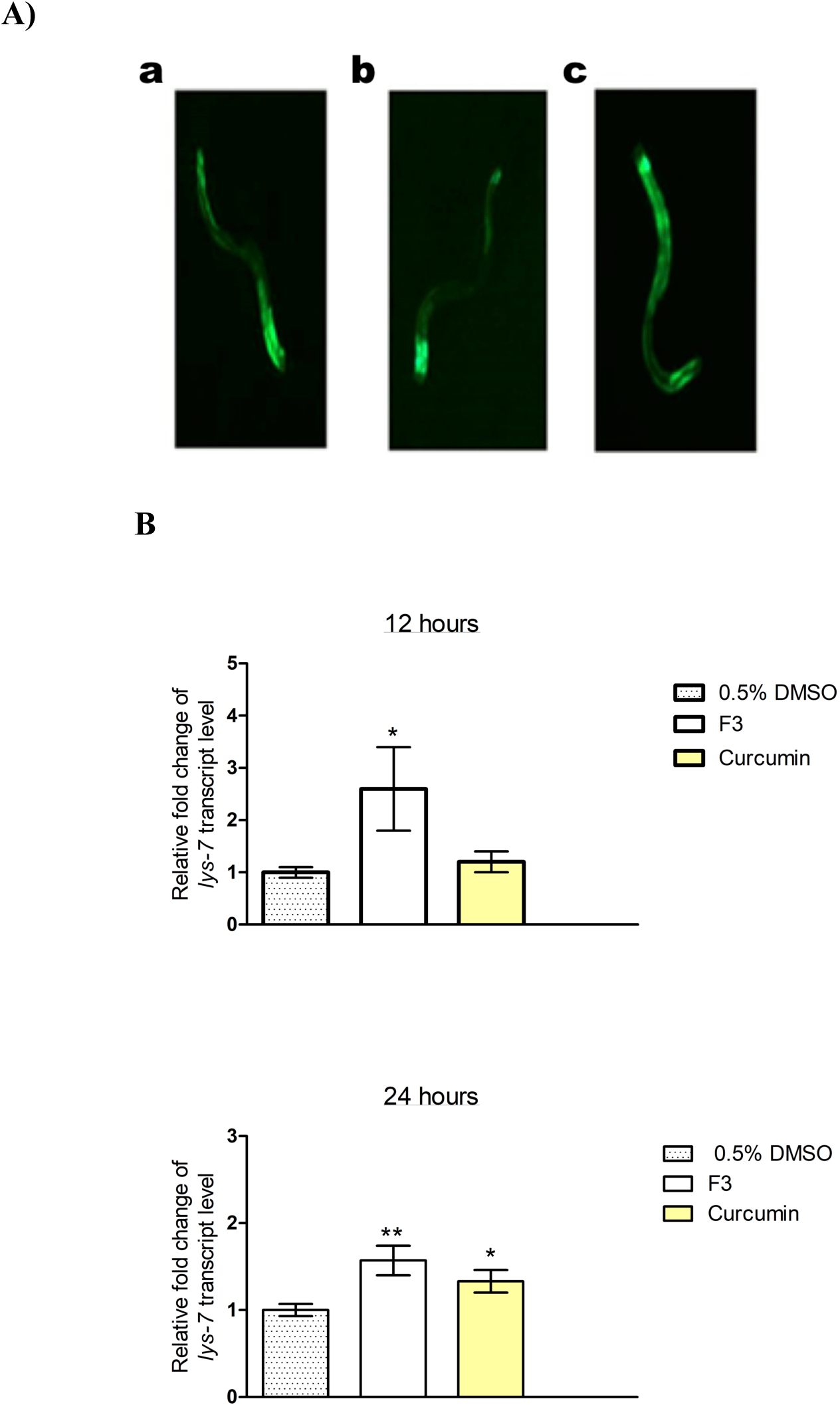
A) *Holothuria atra* restore *lys-7* expression in *P. aeruginosa* PA14 infected *C. elegans*. Representative fluorescence micrographs (60x magnification) of transgenic *lys-7*::GFP worms fed with *E. coli* OP50 (a), infected with *P. aeruginosa* PA14 and treatment with 0.5% DMSO (b), infected with *P. aeruginosa* PA14 and treatment with F3 (c). Images were captured at 12 hr after treatment. (**B**) qPCR of *C. elegans lys-7* in PA14 infected worms treated with 0.5% DMSO, *H. atra* F3 fraction or curcumin at 12 or 24 hrs. Data were analyzed using Student’s *t*-test where * and ** denotes significance at *p*<0.05 and *p*<0.01 respectively when normalized to 0.5% DMSO treatment.

### 3.3 Tentative identification of major compounds in *H. atra* F3 and F4 fractions

QTOF-LCMS separation detected two major peaks in F3 at Retention Times (*R*_T_) 17.764 min and 18.395. The compounds were identified using an HR MS (accurate mass in negative mode) and HR MS/MS spectra (fragmentation) that matched with METLIN, MassBank and Chemspider databases. From the accurate mass analysis (Table 1), the molecular formula was calculated as C_18_H_22_N_2_ and C_10_H_12_N_4_O_5,_ respectively. The compounds were putatively identified as aspidospermatidine and inosine.

**Table 1.** Aspidospermatidine and inosine, the main compounds identified in F3 fraction of *Holothuria atra*

| ESI-MS<br>RT (min) | Detected ion<br>(*m/z*) [M-H]<sup>-</sup> | Fragments | Molecular Formula | Identification |
| --- | --- | --- | --- | --- |
| 17.764 | 266.1789 | 266, 144,<br>136, 130 | C<sub>18</sub>H<sub>22</sub>N<sub>2</sub> | Aspidospermatidine |
| 18.395 | 268.194 | 268, 215,<br>185, 136,<br>133, 109, 93,<br>82, 54 | C<sub>10</sub>H<sub>12</sub>N<sub>4</sub>O<sub>5</sub> | Inosine |

## 3. Experimental

### 3.1 Bacterial and Nematode Strains

*P. aeruginosa* strain PA14^26^, which is rifampicin-resistant, was grown in King’s B broth supplemented with rifampicin (100 µg/mL) while the streptomycin-resistant *E. coli* strain OP50^27^ was cultured in Luria Bertani (LB) broth with streptomycin (100 µg/mL). Cultures were grown overnight with aeration at 37°C. The wild type *C. elegans* Bristol N2 (N2) and transgenic *lys-7*::GFP worm strain SAL105 were obtained from the *Caenorhabditis* Genetics Center (CGC), USA, with growth and handling as described elsewhere.^27^ Standard Operating Procedures involving *C. elegans* and living modified organisms (LMOs) were approved by the Universiti Sains Malaysia Animal Ethic Committee (AECUSM) and the Institutional Biosafety Committee (UKKP)

### 3.2 Preparation of *H. atra* extract

*Holothuria atra* was collected from Pulau Songsong Kedah situated at the northern region of Peninsular Malaysia (N5°48’33.72” E100°17’32.63”) and authenticated by the Centre for Marine and Coastal Studies, Universiti Sains Malaysia. Samples were stored at −20°C until further processing. Frozen samples were cleaned with distilled water, sliced into smaller pieces and freeze-dried for about 5-7 days. Dried samples were extracted with either methanol or acetone, followed by 15 minutes of sonication at room temperature and lastly, filtration with Whatman filter paper No. 1. Extraction procedure was repeated thrice. Filtrates were then concentrated using a rotary evaporator at 39°C. All dried extracts were kept at −20°C before dilution in DMSO at 20mg/mL, followed by storage at 4°C. Depending on assays, the extract stock solution is further diluted to achieve the desired concentrations.

### 3.3 *C. elegans* survival Assay

The slow killing assay was performed according to the protocol described elsewhere.^28^ PA14 cultured overnight on King’s B broth was evenly spread on Pseudomonas Infection Agar (PIA) assay plates supplemented with natural products, followed by incubation at 37°C for 24 hours to allow growth of PA14. Triplicate plates containing 50-60 young adult worms each were used. At least three independent repeats were performed.

Percentage survival of the infected worm population was determined after 48 hours of incubation at 25°C. DMSO (0.5%) and curcumin (100 µg/mL), a known anti-infective agent, was used as negative and positive control, respectively.^29^

### 3.4 Antimicrobial test

The minimum inhibitory concentration (MIC) broth microdilution antimicrobial test was carried out to determine bactericidal activities of natural product. Stock solution of extracts or fractions were further diluted in DMSO to achieve concentration ranging from 25 µg/mL to 1000 µg/mL. Overnight culture of *P. aeruginosa* PA14 was adjusted to 0.5 McFarland standard turbidity and diluted to 1:20. A total of 10 µL of inoculum was added into the wells, followed by incubation at 37°C for 24 hours. Streptomycin (100 µg/mL) and DMSO (0.5%) were also tested as positive and negative controls, respectively.

### 3.5 Bacterial growth assay

The assay was conducted with similar preparations as the MIC broth microdilution test. Upon inoculum addition, the well plate was incubated at 37°C with aeration and bacterial cell density was measured at OD_600_ every 2 hours.

### 3.6 Liquid-liquid partitioning of methanolic *H. atra* extract

The methanol extract was subjected to liquid-liquid chromatography, to separate compounds according to polarity. The extract was suspended in distilled water and subsequently partitioned with butanol. The partition was collected and dried with a rotatory evaporator, followed by storage at −20°C. Stock solution for the partition was prepared in DMSO and stored at 4°C.

### 3.7 Fractionation of *H. atra* butanol partition

The butanol partition was column chromatographed over silica gel 60 (70-230 mesh) and eluted stepwise from 100:0:1 to 0:100:1 with the chloroform:methanol:water solvent system at ratio of 80:20:1. The process was repeated until sufficient fractions were obtained.

### 3.8 Virulence factors and biofilm production assays

Protease and elastase activity assays were performed as described elsewhere, with minor modifications.^30^ Briefly, *P. aeruginosa* PA14 was grown in the presence of bioactive fraction at 200 µg/mL or DMSO (0.5%) for 24 hours at 37°C. The supernatants were collected and filtered with a 0.22 μm nylon filter. Reaction mixture consisting of 0.8% azocasein in 500 µL of 50mM K_2_HPO_4_ (pH7) and 100 µL of the purified supernatant were incubated at 25°C for 3 hours. A total of 0.5 mL of 1.5M HCl was added into the reaction, followed by placement on ice for 30 minutes and centrifugation. After the addition of 0.5 mL 1M NaOH, the absorbance was measured at 400nm (SpectraMax M5).

As for elastase activity, 100 μL of the purified supernatant described above was added into tubes pre-prepared with 1 mL of 10mM Na_2_HPO_4_ and 20 mg of elastin-Congo red. The mixtures were then incubated at 37°C for 4 hours with agitation. After the incubation, the mixtures were centrifuged at 14 000 rpm for 10 minutes and absorbance determined at 495nm.

The pyocyanin quantification assay was conducted as described.^31^ *P. aeruginosa* PA14 was grown in the presence of bioactive fraction for up to 24 hours at 37°C. Cultures were collected at 6, 12 and 24 hours after treatments and centrifuged. A total of 3 mL chloroform was then added into the resulting supernatant, vortexed briefly and centrifuged for 10 minutes at 10 000 rpm. The bottom blue layer was isolated, and the extraction processes were repeated using 1.5 mL of 1M HCl. The absorbance at 520 nm of the pink layer was measured. Pyocyanin concentration was determined by multiplying the absorbance value with 17.072.

A microtiter biofilm formation assay was used to determine the effect of *H. atra* fraction on *P. aeruginosa* PA14 biofilm formation.^32^ *P. aeruginosa* PA14 was grown overnight followed by dilution to 1:100. A total of 100 µL of the inoculum was added into 96 wells plate preseeded with *H. atra* fraction or 0.5% DMSO. The plate was then incubated for 24 hours at 37°C. Liquid and cells were discarded, followed by rinsing of plate with distilled water. For visualisation, 125 μL of 0.1% crystal violet solution was added into each well, followed by incubation for 15 minutes at room temperature. Excessive stain was washed off with water and wells left air-dried for 2 hours. Stained biofilm was treated with 125 μL of 30% acetic acid followed by 10-15 min incubation. A total of 125 μL of solubilized crystal violet was transferred into a new microtiter for measurement at 550 nm.

### 3.9 Visualization of transgenic *lys-7*::GFP *C. elegans*

Transgenic *C. elegans lys-7*::GFP strain SAL105 were exposed *P. aeruginosa* PA14 in the presence or absence of *H. atra* fractions as described in the slow killing assay. The fluorescence intensities of young adult worms at 12 hpi for all the experimental conditions were visualized using Leica Microsystem M205 FA microscope and digitally captured with Leica DFC310 FX camera (60x magnification)

### 3.10 Real-time PCR based quantification of *lys-7* expression in *C. elegans*

Primers were designed accordingly for the amplification of *C. elegans lys-7* and pan-actin as housekeeping gene. Total RNA was treated with RNase-free DNase (Promega, USA) for removal of genomic DNA. PCR optimization and amplification was performed under condition described previously.^28^ Melt curve analysis was performed to ensure the specificity of the PCR products. Quantitative expression of *lys-7* was calculated using CFX Manager software (BioRad, USA) by normalizing its level to housekeeping gene. Three independent replicates were performed for each analysis.

### 3.11 Chemical profiling of fractions

Compounds identification of both the *H. atra* active fractions were performed using the Agilent 6520 accurate-mass quadrupole time-of-flight liquid chromatography mass spectroscopy (QTOF-LCMS). Zorbax SB-C18 column (1.5µm particle size, 0.5×150mm) was used and separation was achieved with a 22.5 minutes gradient of 3-90% acetonitrile, followed by 3% acetonitrile for 10 minutes at a flow rate of 20 µL/min. Solvents contained 0.1% formic acid. The compound prediction was achieved by comparing the obtained individual MS/MS fingerprinting spectrum with METLIN, MassBank and Chemspider databases.

### 3.12 Statistical analysis

Data were analyzed using Student’s *t*-test whereby in all comparisons, *p*<0.05, *p*<0.01 and *p*<0.001 were accepted as statistically significant.

## 4. Discussion

An ongoing strategy to overcome the problem of drug-resistant pathogens is discovery of anti-infective activity, which either disrupt virulence-related pathways or boost host immunity. The diverse marine environment imposes the selection of an array of metabolites produced by an immeasurable source of organisms. Marine natural products with anti-virulence or immunomodulatory properties have been reported.^33-37^. Among edible marine organisms, sea cucumber is regarded as a potent source of novel entities useful for both nutraceutical and pharmaceutical industries. There is an increasing reports of bioactivities from *H. atra*, a widely distributed tropical sea cucumber species. This includes antibacterial, antifungal, anti-proliferative, anti-apoptotic and antioxidant properties.^6, 38-43^ We report here for the first time the anti-infective capacity of *H. atra* extract through an *in vivo*

Our results showed that the *H. atra* methanol extract conferred survival to *P. aeruginosa* PA14-infected *C. elegans* without possessing direct bactericidal activity towards *P. aeruginosa* PA14, ruling out bacteria mortality as reason for improved worm survival. Previously, a methanol extract of *H. atra* was also reported to not exert any bactericidal effect on several pathogenic bacteria species, including *P. aeruginosa*.^44^ Further separation of the *H. atra* methanol extract yielded a fraction with similar bioactive properties. The range of worm survival obtained with this fraction is comparable to levels using a similar slow-killing assay where infected worms were treated with a terrestrial plant seed extract^28^ or curcumin.^29^ These levels however, are lower than a fraction isolated from a tropical marine actinomycete.^45^

In order to better understand the mechanisms responsible for the survival of infected worms, we assessed the effect of *H. atra* towards several *P. aeruginosa* PA14 virulence factors. Although the complete spectrum of the pathogenesis of *C. elegans*-*P. aeruginosa* slow killing assay is not fully understood, the involvement of multiple virulence factors such as pyocanin, pyoverdine, various proteases, elastase, phospholipase and exotoxin is canonical.^25, 26^ Results show *H. atra* lowered both proteolytic and elastolytic activities, and diminished the production of pyocyanin and formation of biofilm, collectively signifying the attenuation of *P. aeruginosa* PA14 virulence. In addition, the collective mitigation of these virulence factors suggests a possible interference of PA14 quorum sensing (QS) by *H. atra* metabolites. QS have been corroborated as a crucial factor responsible for host mortality in the *C. elegans* slow killing assay. Elsewhere, a marine seaweed extract was reported to repress *P. aeruginosa* QS genes, leading to diminished production of virulence factors and survival of infected worms.^36^ Future work should ascertain the capacities of *H. atra* metabolites in interfering with pivotal elements of the *P. aeruginosa* QS pathway.

The bioactive fraction was also able to restore the expression of *lys-7*, which encodes an immune-specific lysozyme which facilitates the clearance of bacterial infection through disruption of bacterial cell wall peptidoglycans.^46^ Two plausible reasons could explain the restitution of *lys-7*. A known characteristic of *P. aeruginosa* infection in *C. elegans* is the suppression of host *lys-7* through the subversion of the DAF-2/DAF-16 insulin-like signaling pathway, leading to the downstream transcriptional suppression of several antimicrobial factors, including *lys-7*.^46-48^ Since the *H. atra* F3 fraction could mutually reduce the production of virulence factors and restore expression of *lys-7* in infected worms, it is possible that the latter observation is a consequence of halting the *P. aeruginosa* virulence activities upstream of the DAF2-DAF16 pathway. This would mimic the effects exerted by a marine brown seaweed extract in overcoming the pathogenicity of PA14 towards *C. elegans*.^35^ The second possible explanation is that *H. atra* also contain compounds capable of directly inducing *lys-7* activities.^45, 49^

Compounds with immunomodulatory activities have been isolated from edible sea cucumber species such as *Cucumaria japonica*, *C. frondosa* and *Stichopus japonicus*.^12-15^ Among these anti-infective molecules, polysaccharides are the most studied group.^15, 50^ In addition, some triterpene glycosides, which are prominent secondary metabolites in sea cucumbers, also possessed immune-boosting effects.^10^ Two examples, cucumariosides and frondoside, were reported to promote cellular immunity through stimulation of phagocystosis and lysosomal activities.^12, 14, 51^ In this study, the main constituents detected in the bioactive *H. atra* fraction are aspidospermatidine and inosine. Aspidospermatidine is an indole alkaloid, a large and growing group of potentially useful marine-based metabolites.^52^ Elsewhere, alkaloids have been identified in *H. atra* extract.^42^ Indole and its derivatives have been demonstrated to repress the *P. aeruginosa* QS system, leading to diminished production of protease, pyocyanin and biofilm.^53-55^ Inosine, a deaminated product of adenosine, has also been highlighted elsewhere as having immunomodulatory and neuroprotective properties.^56-58^

## 5. Conclusions

Using *C. elegans* as host model organism, we propose that *H. atra* has anti-infective properties against *P. aeruginosa*. A bioactive fraction of this sea cucumber species inhibits the production of pathogen virulence factors. Concomitantly, *H. atra* also restored the expression level of host *lys-7*. Collectively, this signifies an *in vivo* anti-infective capacity in *H. atra* which maybe mitigate concerns related with drug resistant pathogens.

## Acknowledgments

We thank Universiti Sains Malaysia for postdoctoral appointment for Su-Anne E. Wan-Ting L. was supported by the MyBrain15 Scholarship from the Ministry of Education, Malaysia. This work was supported by the Malaysia Ministry of Science, Technology and Innovation under the IPHARM Flagship Research Grant ‘Development of Nutraceutical and Pharmaceutical Products for Therapeutic Indications of High Burden to Malaysia’ [Grant Number 304/PBIOLOGI/650869/K105].

---

**Supplementary 1.**
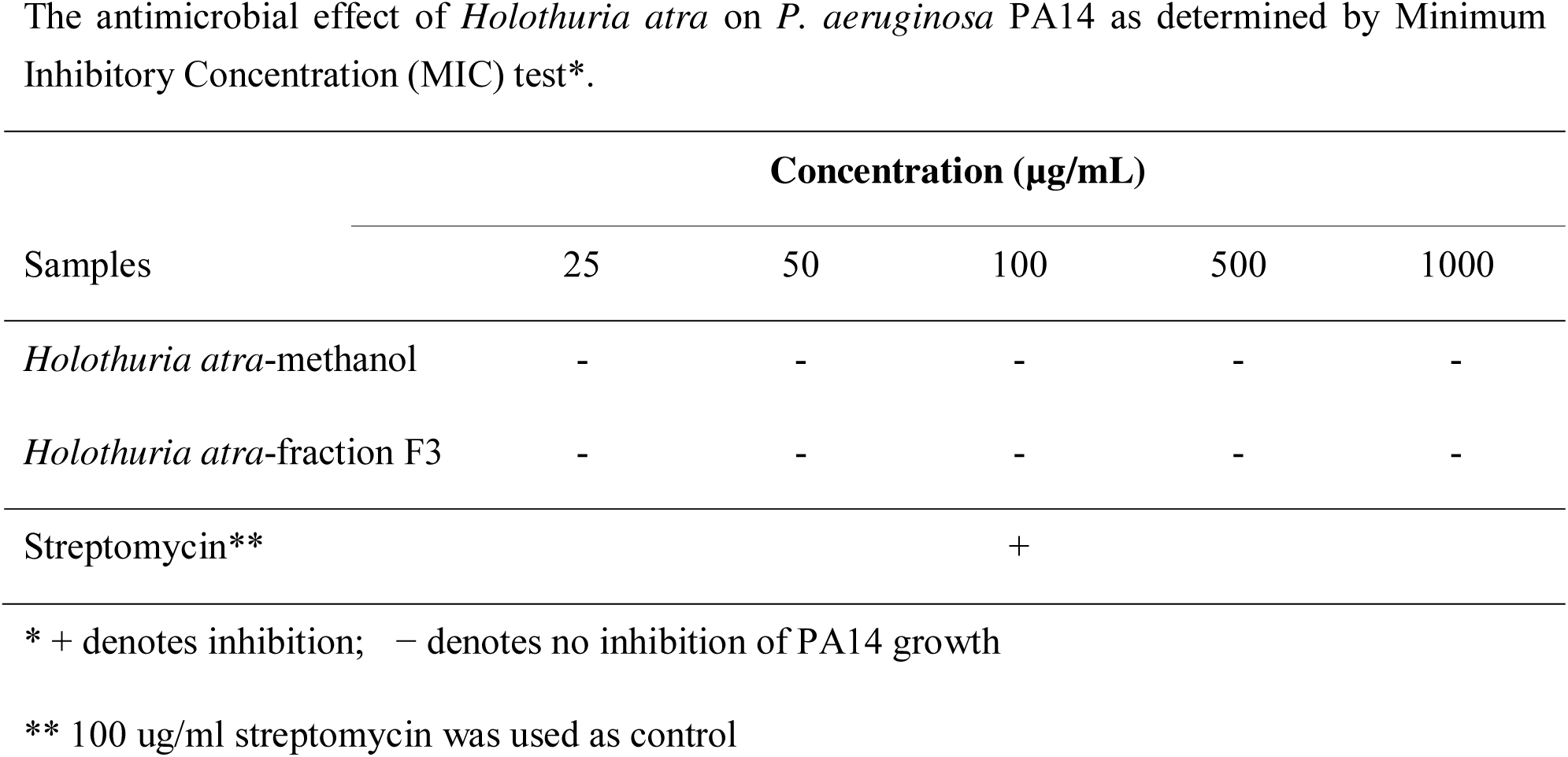
The antimicrobial effect of *Holothuria atra* on *P. aeruginosa* PA14 as determined by Minimum Inhibitory Concentration (MIC) test*.

